# Box-Counting Fractal Dimensions of Cranial Sutures: Effects of Measurement Conditions and Model-Based Reproduction of Fractal-Like Patterns

**DOI:** 10.64898/2026.05.02.722422

**Authors:** Kenta Haishi, Takashi Miura

## Abstract

Cranial sutures are important structures associated with skull growth, and it is widely believed that cranial sutures exhibit fractal structure. However, measurement conditions and analytical procedures have varied among studies, making direct comparison and interpretation difficult. In this study, we reviewed previous measurements and established a standardized box-counting protocol for the fractal dimension (FD) of cranial sutures by quantifying the effects of skeletonization and scale-range selection. Using this protocol, we quantified FD in 45 digitized images of human lambdoid sutures and in eight structure-formation model variants: step, Gaussian, and Mexican-hat kernels; absolute-value and sine *F*_base_ fields; low-frequency noise; a time-dependent kernel; and a given-pattern initialization. We tested whether each model can generate structures with FD values at least as high as those of real sutures using one-sided Welch *t*-tests with Bonferroni correction for eight comparisons (*α*_adj_ = 0.00625), and found that six of eight model variants satisfied this criterion. Analysis of scale-dependent FD further revealed that FD is close to 1 at fine scales and approaches 2 at coarse scales in both real and model-generated patterns, indicating that cranial sutures are better described as finite-range fractal-like structures than as strict fractals, and that the box-counting FD is a scale-conditioned descriptor sensitive to preprocessing choices.

## 1. Introduction

Skull sutures are thin connective tissues that connect skull bones (Drake et al., 2014) and are extensively studied in various academic fields. At birth, human cranial sutures are relatively wide and straight; as the skull grows, they become narrower and increasingly interdigitated, developing the complex morphology characteristic of adult sutures (Fig. 1A). From a clinical point of view, suture tissue serves as a growth center during adolescence, and the premature closure of suture tissue results in craniosynostosis, which requires surgical treatment in severe cases (Cohen & MacLean, 2000). In developmental biology, various growth factors, such as transforming growth factor beta (TGF-*β* ), fibroblast growth factor (FGF), and bone morphogenetic protein (BMP), are known to influence suture patency and interdigitation (Twigg & Wilkie, 2015). It is especially well known that the constitutively active FGF receptor causes craniosynostosis (Cohen & MacLean, 2000; Twigg & Wilkie, 2015). The relationship between genes and pattern formation remains an active area of research.

**Figure 1.**
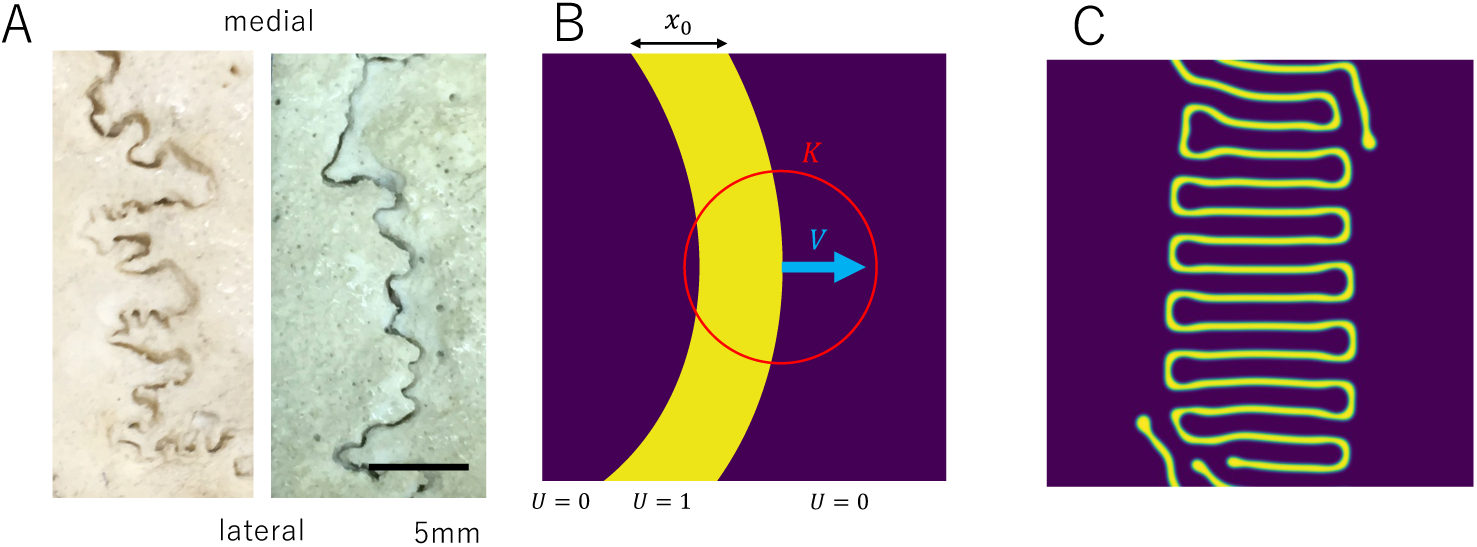
(A) Patterns of adult human lambdoid sutures. (B) Model scheme (Yoshimura et al., 2016). Suture tissue and bone are represented as *U* = 1 and *U* = 0, respectively. The effect of the diffusible signaling molecule is defined by kernel *K*. Suture tissue has width *x*_0_ initially. Interface speed between bone and suture tissue is defined as *V*. (C) Example result of the model using a Gaussian kernel.

Suture morphology is related to its function. For example, Cranial biomechanics influences the shape of suture morphology. As the skull grows and experiences functional loads such as mastication, suture interdigitation develops in relation to local mechanical demands. Interdigitation enables collagen fiber arrangements that resist deformation, particularly under compression, helping explain its prominence in mechanically loaded regions (Herring, 2008). Cranial suture morphology varies among mammalian species and is associated with factors such as diet and behavior (Buezas et al., 2017; Byron, 2009). This biological variation should be considered when comparing fractal-dimension measurements across studies involving different species, including non-human taxa (White et al., 2020; 2026).

Skull suture has been reported to be a fractal structure. A fractal structure is a geometrically complex pattern that retains similar irregularity across spatial scales, and its fractal dimension (FD) quantifies how densely it fills space. Two widely used methods for estimating fractal dimension are the box-counting and divider methods. The box-counting method quantifies how the number of boxes required to cover a structure scales with box size, whereas the divider method quantifies how the measured length of a curve scales with the ruler length used to trace it. The first report on the fractal nature of sutures appeared in 1985 (Long, 1985). They showed that the complex sutures of deer skulls and ammonites are extremely long relative to the areas they enclose, repeat similar details at several scales, and have fractal dimensions between 1 and 2. Masuda and Yohro found that human sagittal, coronal, and lambdoid sutures are fractal curves with dimensions between 1.2 and 1.5, that this dimension does not differ by suture, age, or individual, and that the wavelength of the sagittal suture changes across the growth, fusion, and obliteration stages. (Masuda & Yohro, 1987).

Subsequently, numerous measurements of the fractal dimension of sutures were conducted. Their purpose varies: assessment of the measurement method (Hartwig, 1991; Long & Long, 1992; Górski & Skrzat, 2006; White et al., 2020), description of human suture shape (Yu et al., 2003; Skrzat et al., 2004), relationship with age or maturation (Lynnerup & Jacobsen, 2003; Darawsheh et al., 2023; Costamagna et al., 2026), Explaining differences among species, regions, and functions (Gibert & Palmqvist, 1995; Monteiro & Lessa, 2000; Nicolay & Vaders, 2006; Buezas et al., 2017; White et al., 2026; Di Guida & Cassini, 2026), and understanding pattern formation mechanisms (Miura et al., 2009). Some studies validate the box-counting method by measuring theoretical fractals whose dimensions are known. (Costamagna et al., 2026; & Cassini, 2026). Currently, the fractal dimension is one of the most frequently used quantitative measures for describing the complexity of the structure. However, there exist some discrepancy of FD between studies. For example, Górski & Skrzat (2006) estimated FD of sagittal suture as 1.56 while (2009) estimated it about 1.2. Because image resolution, preprocessing, and box-size range can affect FD estimates, a standardized box-counting protocol is needed to facilitate meaningful comparisons across studies.

Various models have been proposed to explain and generate the fractal structure of skull sutures. Some models incorporate the collision of two randomly growing rough interfaces (Eden, 1961; Oota et al., 2004; 2006) or diffusion-limited aggregation to generate a fractal suture structure (Zollikofer & Weissmann, 2011). These approaches represent the suture as a widthless interface and do not explicitly address craniosynostosis. Other models utilize skull osteogenesis and a diffusible signaling molecule to reproduce suture width maintenance and interdigitation (Miura et al., 2009; Yoshimura et al., 2016). Further studies explored whether a time-dependent diffusion coefficient (Miura et al., 2009) or spatiotemporal noise (Naroda et al., 2020) could reproduce the fractal properties of sutures. However, the mechanism for fractal structure generation is only partially understood.

In the present study, we established a standardized box-counting protocol by quantifying the effects of skeletonization and scale-range selection, and used it to compare human lambdoid sutures with mathematical model variants to identify candidate mechanisms capable of generating FD values at least as high as those observed in real sutures. First, we surveyed previous FD studies and quantified how differences in preprocessing, especially skeletonization (reducing a shape to a one-pixel-wide centerline while preserving its over-all topology), and in the selected box-counting scale range can affect FD estimates and obscure direct comparisons among previously reported FD values. Second, we applied this protocol to human lambdoid sutures and to model-generated patterns to test which candidate pattern-formation mechanisms can generate FD values equal to or exceeding those of real sutures.

## 2. Materials and Methods

### 2.1 Literature search

To summarize previous reports on fractal dimension (FD) measurement of cranial sutures, we performed a targeted literature search in PubMed using the keyword string “suture fractal”. Retrieved articles were screened manually, and studies not focused on biological or anatomical specimens were excluded. Because the aim of this review was to summarize methodological variation, studies from non-human species were retained. From the eligible studies, we extracted the target sample, structure, age, FD measurement method, software, reported FD values, and whether skeletonization was applied. This search was intended to provide a narrative overview of methodological variation rather than a formal systematic review or meta-analysis.

### 2.2 Measurement of fractal dimension

We used 45 two-dimensional images of human lambdoid sutures (512 × 512 pixels); specimens were from Kyushu University School of Medicine and School of Dentistry. Imaging conditions and image-processing procedures are described in our previous study (Naroda et al., 2020). Each image was binarized and skeletonized before FD measurement. Real suture images were binarized using Otsu’s automatic threshold (Otsu, 1979); model outputs were binarized at a fixed threshold of *u >* 0.5. Skeletonization was applied to both using `skimage.morphology.skeletonize`. This work was approved by Kyushu University Institutional Review Board for Clinical Research (2019-350).

We used the box-counting method to measure the fractal dimension. The relationship between box size and suture pattern was visualized to determine the measurement scale range. The software to specify the scale range and calculate box-counting FD is available via GitHub (https://github.com/haiken-hub/ScalingFractalTools).

Box sizes were [1.2*^k^*] pixels for integer *k* = 0, 1,. 25. Largest box size is 95, which corresponds to 19% of the image size. Larger box sizes results in same box count in different box size, resulting in stair-like distribution which may affect regression. Skeletonization reduced each suture to a one-pixel-wide line, equal to the minimum box size, so the line width was not counted across multiple boxes. When rounding produced duplicate integer box sizes, box counts at identical sizes were averaged before regression. Fractal dimension was estimated as the negative slope of log *N* vs. log *s* over *k* = 0,…, 25 (26 box sizes) for the primary model–real comparison. FD computation used the `box_scaler_preview` function from ScalingFractalTools.

### 2.3 Base model

We chose a preexisting suture pattern formation model (Yoshimura et al., 2016) (Fig. 1B, C). In this model, the mesenchymal tissue secretes growth factors that have osteogenic ability within a certain range determined by a predefined function (kernel). The function is point-symmetric because the effect of the diffusible signaling molecule does not have a preferred direction of diffusion. We also assume that the osteogenic front moves according to the effect of the diffusible signaling molecule. In such a system, a slight initial perturbation of shape is amplified while maintaining suture width, resulting in the formation of the interdigitation pattern (Fig. 1C).

### 2.4 Models of skull suture pattern formation

Next, we aimed to reproduce fractal-like suture patterns *in silico* by introducing biologically plausible modifications into an established modeling framework. We used a base model in which bone differentiation and dedifferentiation depend on the local concentration of growth factor secreted from suture tissue (Materials and Methods). We modified this model in three ways: spatial pattern of growth-factor effects, inhomogeneity in osteogenic differentiation tendency, and stage-dependent changes in ossification properties. For each variant, we ran simulations on 512 × 512 two-dimensional lattices and obtained 45 synthetic suture patterns as output images. We then estimated FD of these patterns using the standard box-counting and skeletonization procedures described above.

#### Spatial pattern of growth-factor-mediated interactions (kernel)

First, we systematically examined whether the *spatial range* of growth-factor-mediated interactions affects the fractal dimension of the simulated suture pattern. In our model (Materials and Methods), each lattice site exchanges signals with others over a finite neighborhood; the weight assigned to neighbors at distance *r* is specified by a function *K*(*r*), which we refer to as the *interaction kernel*. *K* describes how strongly local tissue states at one location influence differentiation at another. We compared three kernel shapes (Fig. 4): a step kernel (uniform influence within a fixed radius), a single-scale Gaussian kernel (one characteristic length *l*_1_), and a Mexican-hat-type kernel implemented as the difference of two Gaussians (two characteristic lengths *l*_1_ and *l*_2_). We hypothesized that kernel shape may influence the complexity of the resulting pattern, so we tested several kernel shapes. For each kernel, we ran otherwise identical simulations on 512 × 512 lattices and estimated the fractal dimension of the resulting patterns using the box-counting procedures described above.

##### Step Kernel

The step kernel assumes that spatial interaction is constant up to a certain length. This kernel is used in the original model (Yoshimura et al., 2016).

##### Gaussian Kernel

The Gaussian kernel assumes that spatial interaction decreases smoothly with distance (Fig. 4).

##### Mexican Hat Kernel

We also used the Mexican hat kernel, assuming short-range stimulation and long-range inhibition. We expected that a more complex kernel shape might result in more complex structures.

#### Inhomogeneity in osteogenic differentiation tendency

##### Regional difference

To keep the interdigitating suture within a certain width, as observed in the skull, we implemented a bone differentiation promotion factor as *F*_base_, which is designed to restrict the expansion of suture tissue depending on the distance from the center of the pattern formation region (Fig. 5). By using this function, the winding suture is kept within a certain width even after long simulation times. We expect that if the pattern formation parameter differs between the center and periphery of the field, it can create regional differences in the pattern, resulting in increased complexity.

##### Low-frequency noise

In addition, we added low-frequency noise (Fig. 5C) to increase the complexity of the resulting structure. It is known that white noise plus a smoothing effect can produce a structure that shows fractality (Naroda et al., 2020).

#### Time-dependent parameter

##### Time-dependent kernel

We used a time-dependent kernel shape. We assumed that two characteristic lengths in the early and late stages of development may lead to a scaling property (Miura et al., 2009).

##### Approximation of early-stage pattern (given-pattern model; GP)

As an alternative to running the full early-stage simulation, we approximated the early-stage pattern as a sine curve and used it as the initial condition for the late-stage kernel (Supporting Information Section S3). “GP” here refers to the *given-pattern* initialization strategy and is unrelated to Gaussian processes in the statistical sense.

### 2.5 Numerical simulations

All numerical simulations were implemented in Python (Van Rossum & Drake, 2009). Parameter values used in all simulations are listed in Table 2. Further details of the model are provided in Supporting Information Section S3, and the source code is available on GitHub (https://github.com/haiken-hub/SutureModel).

The same FD-estimation protocol was then applied to both real and simulated patterns so that model–data comparison was not confounded by differences in preprocessing or scale selection.

### 2.6 Statistical analysis

The primary question was whether each model variant can generate structures with a mean FD at least as high as that of real sutures (*H*_0_ : *µ*_model_ *↔ µ*_real_; *H*_1_ : *µ*_model_ *> µ*_real_). We therefore used one-sided Welch *t*-tests, which do not assume equal variance and are appropriate for comparing independent groups of unequal precision. To account for multiple comparisons across eight model variants, Bonferroni correction was applied (*α*_adj_ = 0.05*/*8 = 0.00625). For the scale-dependence analysis, paired *t*-tests were used because each image was measured under both conditions. All analyses were performed in Python using `scipy.stats`.

## 3. Results

### 3.1 Narrative review of previous suture FD measurement

We identified 19 studies that measured the fractal dimension (FD) of suture lines (Table 1). The reviewed studies differed not only in measurement method and preprocessing, but also in biological context, including species, suture identity, and age or developmental stage where reported.

**Table 1.**
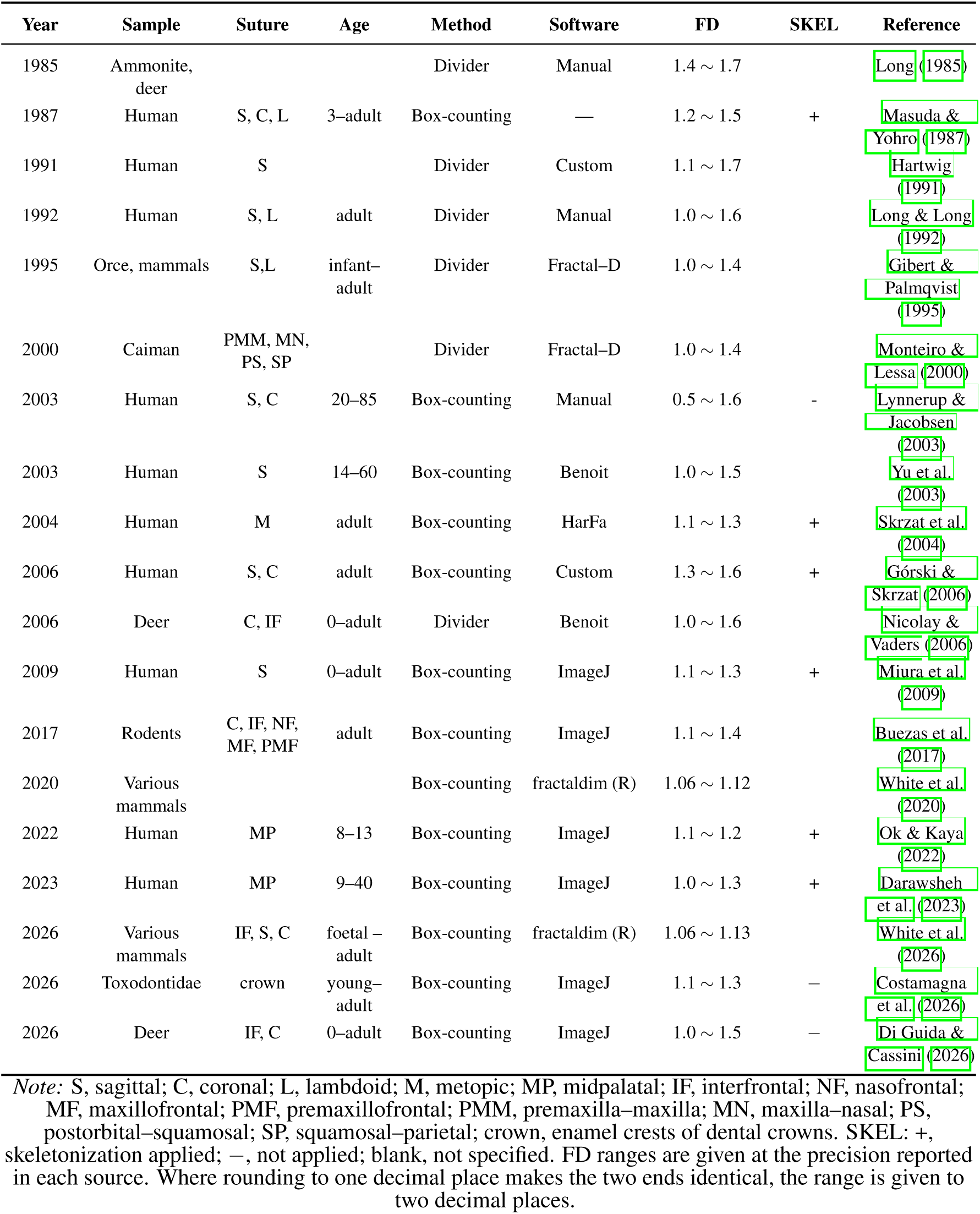
Previous studies measuring fractal dimensions of cranial sutures.

**Table 2.**
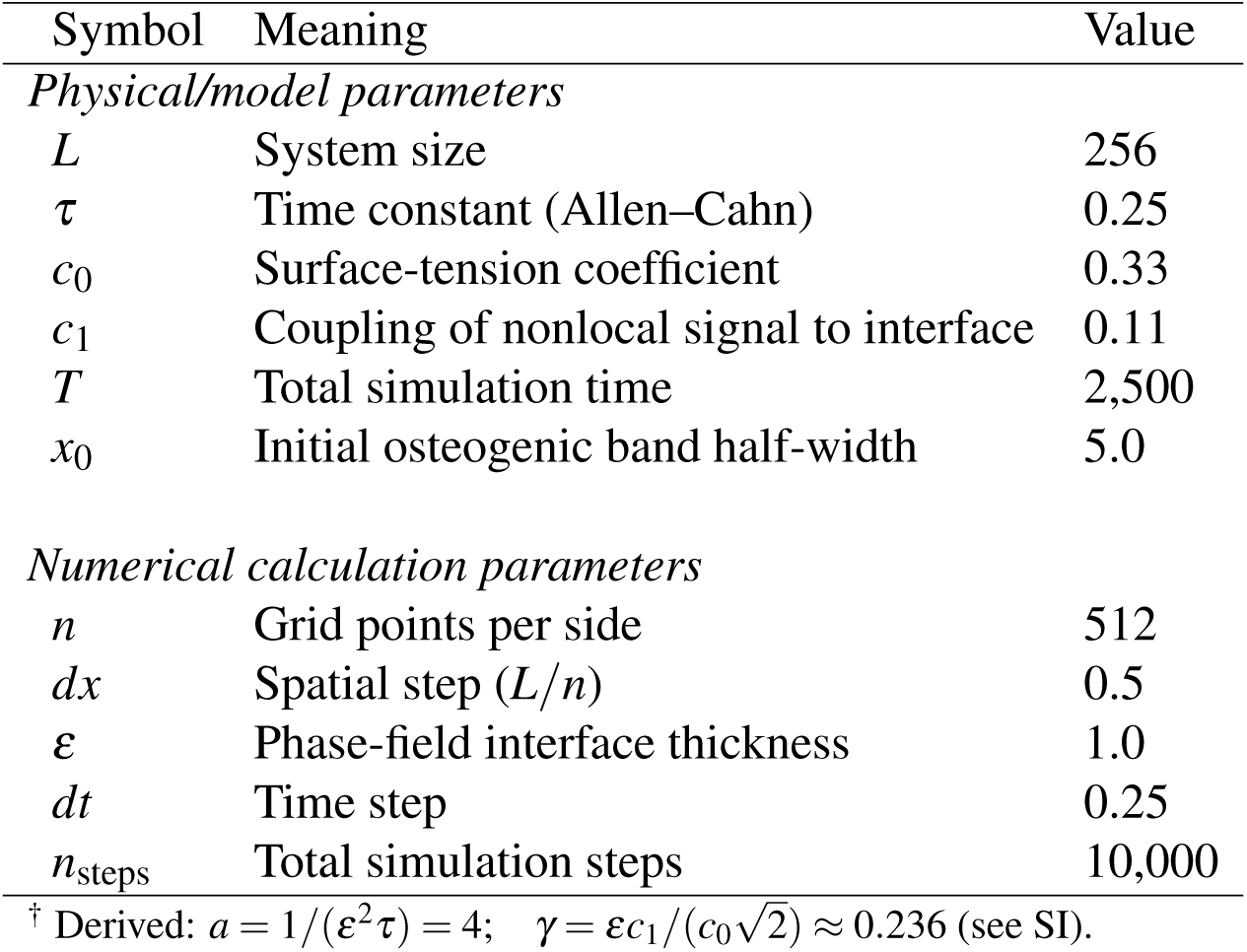
Shared model and numerical parameters used in all simulations.

| Symbol | Meaning | Value |
|---|---|---|
| *Physical/model parameters* |  |  |
| *L* | System size | 256 |
| *τ* | Time constant (Allen–Cahn) | 0.25 |
| *c*<sub>0</sub> | Surface-tension coefficient | 0.33 |
| *c*<sub>1</sub> | Coupling of nonlocal signal to interface | 0.11 |
| *T* | Total simulation time | 2,500 |
| *x*<sub>0</sub> | Initial osteogenic band half-width | 5.0 |
| *Numerical calculation parameters* |  |  |
| *n* | Grid points per side | 512 |
| *dx* | Spatial step (*L*/*n*) | 0.5 |
| *ε* | Phase-field interface thickness | 1.0 |
| *dt* | Time step | 0.25 |
| *n*<sub>steps</sub> | Total simulation steps | 10,000 |

<sup>†</sup> Derived: *a* = 1/(*ε*<sup>2</sup>*τ*) = 4; *γ* = *εc*<sub>1</sub>/(*c*<sub>0</sub>√2) ≈ 0.236 (see SI).

Concerning the measurement method, there are two major methods of FD estimation. The divider method estimates fractal dimension by measuring how the apparent length of a boundary changes with the length of the measuring step, whereas the box-counting method estimates it from how the number of boxes required to cover the structure changes with box size. The box-counting method was used in the largest number of reports (13), followed by the divider method (6). By year, the divider method was used relatively more frequently in reports prior to 2003, whereas the box-counting method was predominant in reports from 2003 onward. The software used varied widely, including manual measurement, Benoit, HarFa, ImageJ, and fractaldim(R); even for the same method, the implementation environment was not standardized. Two recent studies stated the box-size range. (2026) measured the fractal dimension of enamel crests on toxodont dental crowns, using box sizes from 5 to 1028 pixels. measured interfrontal and coronal sutures of Neotropical deer, with a minimum box size of 5 pixels and a maximum of 45% of the foreground pixels.

There were six reports that described skeletonization (SKEL, Fig. 2A, B) as part of image preprocessing, accounting for approximately 32% of the total. On the other hand, there were also several reports that did not specify whether skeletonization was performed, and the description of preprocessing conditions was inconsistent. Most reported FD values were between about 1.0 and 1.6, often near 1.2. One study fell outside the interval expected for an interdigitated curve (1 *<* FD *<* 2): reported values down to 0.5 (Table 1).

**Figure 2.**
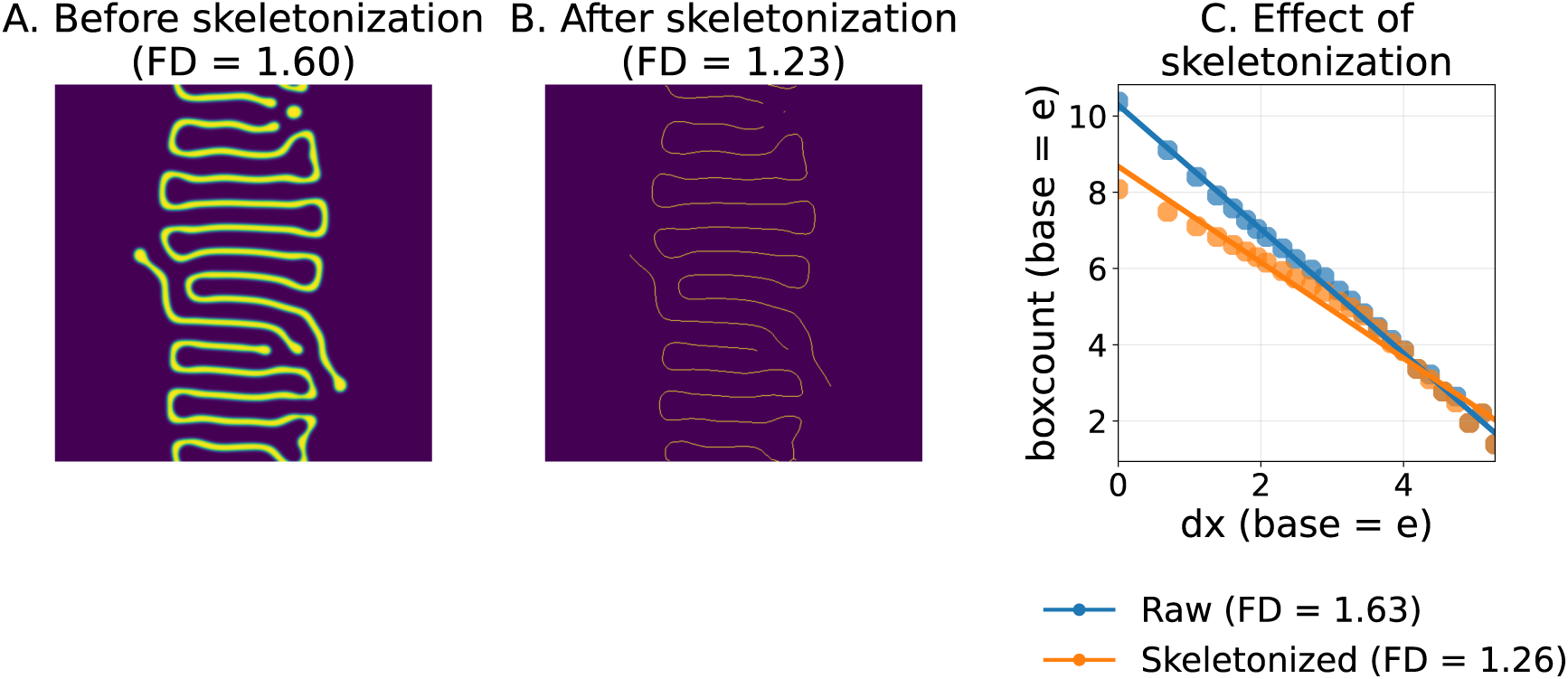
Effect of skeletonization. (A) Before skeletonization. (B) After skeletonization. Suture width becomes irrelevant and the pattern becomes more one-dimensional. (C) Fractal dimension before and after skeletonization.

In summary, while the box-counting method has become the mainstream approach in recent literature, there has been no consensus across studies regarding image preprocessing conditions—including whether skeletonization is applied—or the criteria for scale selection in the box-counting method. These differences in preprocessing were identified as factors that could contribute to variability in FD estimates.

### 3.2 Verification of measurement criteria of fractal dimension

#### Skeletonization

We compared the effect of skeletonization on FD values using a 512 × 512 PNG image generated with a Gaussian-kernel model (Fig. 2C). We estimated the FD by log–log regression of box counts in three scale ranges (all scales: *k* = 0,…, 29; small scales: *dx < e*^3.5^, *k* = 0,…, 19; large scales: *dx ≥ e*^3.5^, *k* = 20,…, 29; base = 1.2), using ten independent Gaussian-kernel realizations (*n* = 10; Table 3).

**Table 3.**
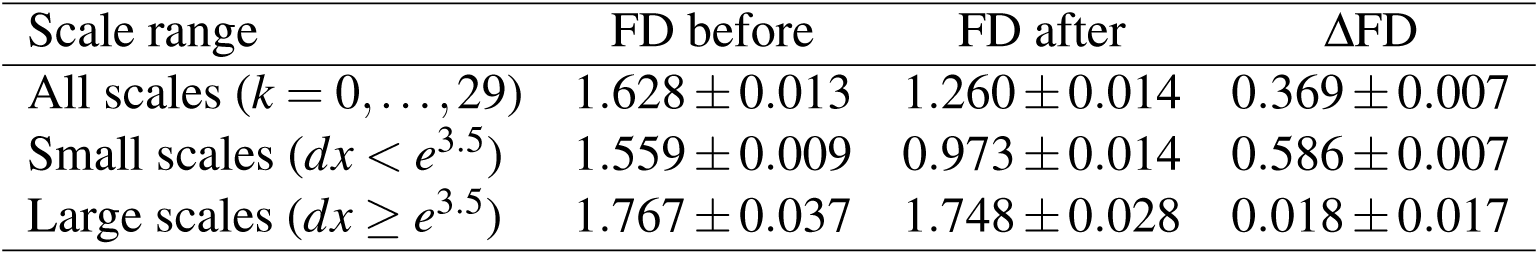
Scale-dependent effect of skeletonization on FD for the Gaussian-kernel model (*n* = 10). FD was estimated by log–log regression of box counts with base 1.2. Small scales: *dx < e*^3.5^ (*k* = 0,…, 19); large scales: *dx ≥ e*^3.5^ (*k* = 20,…, 29); all scales: *k* = 0,…, 29.

| Scale range | FD before | FD after | ΔFD |
|---|---|---|---|
| All scales (*k* = 0, . . . , 29) | 1.628 ± 0.013 | 1.260 ± 0.014 | 0.369 ± 0.007 |
| Small scales (*dx* < *e*<sup>3.5</sup>) | 1.559 ± 0.009 | 0.973 ± 0.014 | 0.586 ± 0.007 |
| Large scales (*dx* ≥ *e*<sup>3.5</sup>) | 1.767 ± 0.037 | 1.748 ± 0.028 | 0.018 ± 0.017 |

Over all scales, skeletonization reduced FD from 1.628 *±* 0.013 to 1.260 *±* 0.014 (!FD = 0.369 *±* 0.007; paired *t*-test, *p <* 10*^-^*^16^). The decrease was most pronounced at small scales (1.559 *±* 0.009 to 0.973 *±* 0.014; ΔFD = 0.586 *±* 0.007; *p <* 10*^-^*^18^), whereas at large scales the change was much smaller (1.767 *±* 0.037 to 1.748 *±* 0.028; ΔFD = 0.018 *±* 0.017; *p* = 7.1 *-* 10*^-^*^3^). Thus, the effect of skeletonization on FD was scale-dependent: large at fine scales and comparatively small at coarse scales. Note that FD values computed on disjoint scale subsets are not directly comparable in magnitude, but the paired before-after differences within each range quantify the scale-dependent effect of skeletonization.

#### Measurement scale

Figure 3A shows the suture line image used for FD measurement, and Figure 3B shows the coverage process at representative scales in the box-counting method. As the scale became coarser, the image was displayed at a larger magnification, making it visually difficult to distinguish the local structures contained in the original image. In particular, at box sizes *⎣*1.2^30^*⎦* and larger (*k ≥* 30), structural information was virtually lost and values usable for measurement were no longer available. Furthermore, the original structure was no longer clearly preserved in the range *k* = 17–21 (Fig. 3B).

**Figure 3.**
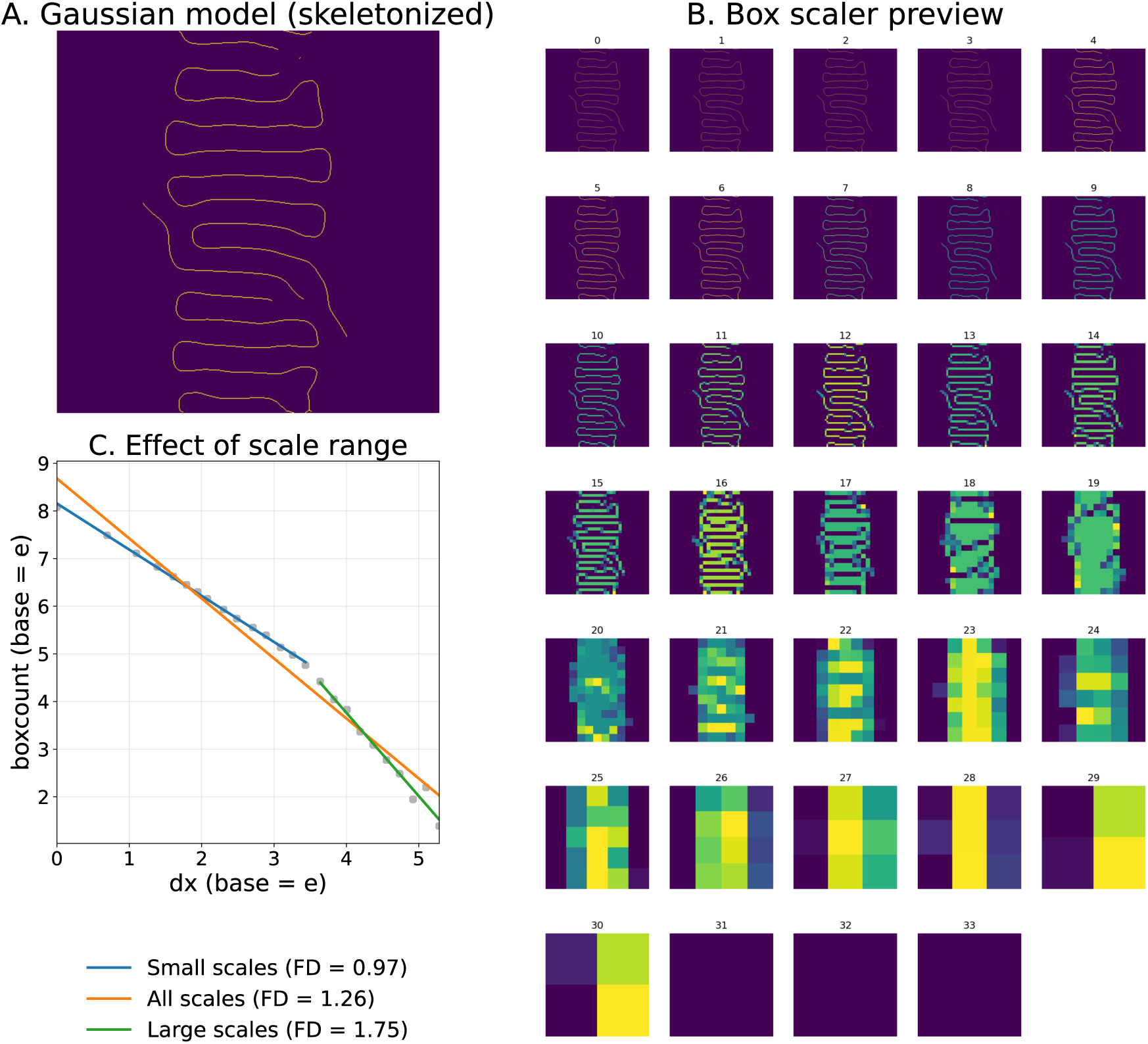
(A) Analyzed model-generated pattern. (B) The relationship between FD and spatial scale. In regions with small box sizes, the pattern is simple and one-dimensional, but as the box size increases, it becomes more two-dimensional. (C) Effect of the selected scale range. The box-counting dimension is higher when the measurement box is large than when it is small.

Based on these observations, we examined the effect of scale selection on FD estimation using 512 × 512 PNG images generated by a Gaussian-kernel model. We defined three index ranges: (i) *k* = 0,…, 29 (full usable range), (ii) *k* = 0,…, 19 (fine scales), and (iii) *k* = 20,…, 29 (coarse scales, where structure begins to be lost). This segmentation aligned with the representative scale at which the log–log slope changes (around *k* = 19, box size *≃* 1.2^19^), as shown in Figure 3C. Regression results showed a slope of 1.0 for *k* = 0,…, 19, 1.7 for *k* = 20,…, 29, and 1.23 for the entire range *k* = 0,…, 29, indicating that the FD estimates are strongly dependent on the chosen scale, consistent with earlier findings that the effective scaling regime is limited by finite resolution and restricted ranges of self-similar behavior (Yu et al., 2003; Long & Long, 1992).

### 3.3 Fractal dimension of patterns generated by mathematical models

The models generated various patterns with FD values larger than or comparable to those of real sutures (Fig. 6-7; model parameters in Supporting Information Section S3). All FD values were estimated over *k* = 0,…, 25 (26 box sizes, base 1.2). The step kernel yielded the highest mean (1.250 *±* 0.011; Δ = +0.152), followed by the given-pattern (GP) dual-stage model (1.172 *±* 0.008; Δ = +0.074) and the time-dependent-kernel (TDK) dual-stage model (1.163 *±* 0.007; Δ = +0.065). The Gaussian kernel (1.149 *±* 0.011; Δ = +0.051) and the ABS parameter field (1.136 *±* 0.011; Δ = +0.038) also lay above the real mean. The Mexican-hat kernel (1.123 *±* 0.014; Δ = +0.025), the sin parameter field (1.128 *±* 0.009; Δ = +0.029), and the low-pass noise parameter field (1.093 *±* 0.013; Δ = *-*0.006) approached the real mean more closely, with the low-pass model showing the best agreement.

Real specimens exhibited a mean FD of 1.098 *±* 0.072 , with values ranging from 0.92 to 1.26, indicating substantial sample-to-sample variability. The lower end of this range falls below 1, outside the interval expected for an interdigitated curve (1 *<* FD *<* 2). In contrast, all numerical models produced much narrower distributions, with standard deviations of only 0.007–0.014—roughly five to ten times smaller than that of the real data (Fig. 7).

To test whether each model can generate structures at least as complex as real sutures in terms of mean FD, we applied a one-sided Welch *t*-test (*H*_0_ : *µ*_model_ *≤ µ*_real_; *H*_1_ : *µ*_model_ *> µ*_real_) for each of the eight models (*n* = 45 per group). At *α* = 0.05, seven of eight models yielded a significantly higher mean FD than real sutures; after Bonferroni correction for eight comparisons (*α*_adj_ = 0.00625), six models remained significant (Table 4). The low-pass noise parameter field did not exceed the real mean (Δ = *-*0.006; *p* = 0.688). The Mexican-hat kernel showed a modest but nominally significant elevation (Δ = +0.025; *p* = 0.013) that became unsupported after Bonferroni correction. The step kernel, dual-stage models, Gaussian kernel, ABS parameter field, and sin parameter field all demonstrated statistically greater mean FD than real sutures under both uncorrected and Bonferroni-adjusted criteria. None of the models reproduced the large standard deviation observed in real sutures.

**Table 4.** One-sided Welch *t*-tests comparing each model with real sutures (*n* = 45). FD was estimated over *k* = 0,…, 25 (26 box sizes, base 1.2) for all groups. Mean difference is defined as (model mean real mean). *H*_0_ : *µ*_model_ *µ*_real_; *H*_1_ : *µ*_model_ *> µ*_real_. Bonferroni-adjusted significance uses *α*_adj_ = 0.05*/*8 = 0.00625.

| Model | Mean diff. | *t* | *p* (one-sided) | 95% CI | Sig. (Bonf.) |
|---|---|---|---|---|---|
| Kernel_Step | +0.152 | 13.81 | < 0.001 | [+0.134, +0.171] | Yes |
| DualStage_GP | +0.074 | 6.72 | < 0.001 | [+0.055, +0.092] | Yes |
| DualStage_TDK | +0.065 | 5.94 | < 0.001 | [+0.047, +0.083] | Yes |
| Kernel_Gaussian | +0.051 | 4.65 | < 0.001 | [+0.033, +0.070] | Yes |
| Parameter_ABS | +0.038 | 3.46 | < 0.001 | [+0.020, +0.057] | Yes |
| Parameter_Sin | +0.029 | 2.68 | 0.005 | [+0.011, +0.048] | Yes |
| Kernel_MexicanHat | +0.025 | 2.28 | 0.013 | [+0.007, +0.044] | No |
| Parameter_LowPass | −0.006 | −0.49 | 0.688 | [−0.024, +0.013] | No |

Figure 9 summarizes the scale dependence of the estimated FD for skeletonized patterns from eight mathematical models and 45 real suture images (*n* = 45 per group). For each sample, FD was obtained independently by log–log regression of box counts (base 1.2) over two scale ranges separated at *dx* = *e*^3.5^: small scales (*dx < e*^3.5^; *k* = 0,…, 19; left panel) and large scales (*dx ≥ e*^3.5^; *k* = 20,…, 29; right panel). Across all 405 samples, the mean FD was 0.98 *±* 0.10 on small scales and 1.68 *±* 0.10 on large scales (Δ = *-*0.70; paired one-sided *t*-test, *p <* 0.001), indicating that FD is systematically lower at fine scales than at coarse scales. This trend was shared by every model and by real sutures (real: FD_small_ = 0.998, FD_large_ = 1.554), although real specimens exhibited substantially broader sample-to-sample variation—particularly in the large-scale regime—whereas numerical models produced tightly clustered estimates. Thus, the box plots visualize a consistent crossover in local FD with measurement scale, rather than a uniform fractal scaling over the full range of box sizes.

## 4. Discussion

### 4.1 Variety of measurement methods in previous work

A comprehensive review of previous reports on the measurement of the fractal dimension (FD) of suture lines revealed some variation in the measurement methods used (Table 1). Representative methods include the divider method and the box-counting method. Theoretically, when the target structure exhibits perfect self-similarity, the dimensions obtained by either method are expected to be the Hausdorff dimension (Supporting Information, Section S1; Falconer, 2003). However, it has been reported that the divider method and the box-counting method may yield different dimensions, even for the same pattern, due to finite image resolution and the influence of noise (Klinkenberg, 1994; Dutch, 1993; Yu et al., 2003). Therefore, the range of FD values summarized in Table should not be interpreted as biological variation in human sutures alone; it reflects a mixture of anatomical, taxonomic, developmental, and methodological differences. Values below 1 or above 2 are not expected for a curve lying between a line (FD = 1) and a filled plane (FD = 2). The entries outside this interval are therefore more plausibly attributed to measurement conditions than to suture geometry. (2003) expected that the fractal dimension decreases to 0 as the suture obliterates and becomes invisible.

For anatomical comparisons and secondary synthesis of published FD values, this finding has an important practical consequence: FD values should be interpreted with caution when the image-processing workflow, skeletonization status (Fig. 2), and scale range (Fig. 3) are not comparable or are not reported in sufficient detail. This is particularly relevant when FD is reused as a compact descriptor of morphological complexity across studies or disciplines. In other words, the fractal dimension is considered an indicator that depends not only on the properties of the structure itself but also on image processing and measurement conditions. The dependence of the fractal dimension on scale selection has been described previously (Long & Long, 1992). Within the interval expected for an interdigitated curve, small numerical differences, such as 1.49 versus 1.54, should not be over-interpreted. Image analysis has a margin of error, and the published values are used to reflect observable differences in suture interdigitation.

### 4.2 Measurement method and FD

When using box-counting for FD measurement, we confirmed that the FD value varies depending on skeletonization and the selected scale. For evaluating FD, we adopted the following measurement conditions: (1) images with skeletonization applied; (2) indices *k* = 0,…, 25 (26 box sizes, base 1.2) for the primary model–real comparison. Sensitivity analyses comparing skeletonization effects and scale-range selection used the broader range *k* = 0,…, 29. This sensitivity is consistent with previous reports showing that FD estimates depend on preprocessing and the selected scale interval (Klinkenberg, 1994; Dutch, 1993; Yu et al., 2003). For these 512 *×* 512 images, the analyses support a largest box of about 19% of the image side (*k* = 0,…, 25), rather than a fixed number of box sizes.

### 4.3 Box-counting dimension of the patterns generated by mathematical models

In this study, we constructed multiple mathematical models to generate patterns with higher box-counting dimensions. Having established that FD estimates depend on pre-processing and scale range, we used the standardized protocol as a common basis for comparing real and model-generated suture patterns. Our original design aimed to introduce a multiscale structure through kernel shape (Fig. 4), boundary conditions, and time-dependent parameters (Fig. 5). Most of the models are capable of generating patterns with FD values equal to or greater than those of real sutures (Fig. 7). This comparison uses the box-counting concerns raised for earlier suture measurements (Hartwig, 1991; Long, 1992; Górski & Skrzat, 2006; White et al., 2020). With preprocessing and scale range held fixed, variants of the osteogenesis model cited in the Introduction reach the FD of real sutures (Miura et al., 2009; Yoshimura et al., 2016). Natural sutures have the highest standard deviation because the specimens differ, whereas each model used one shared initial condition and so varied little across replicates (Fig. 7).

**Figure 4.**
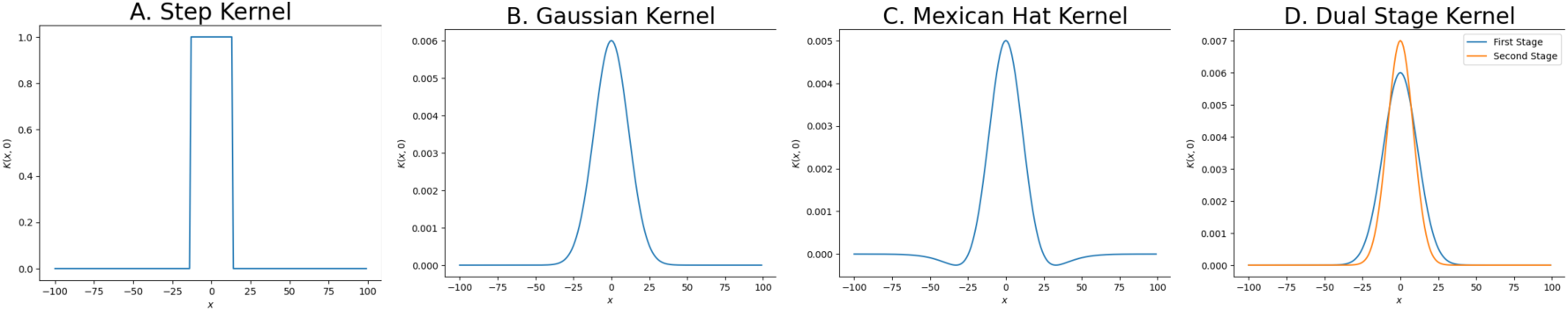
Kernel shapes of the model. We use a point-symmetric function *K*(*x, y*), and the section at *y* = 0 is shown. Four types of kernels are used: (A) step function, (B) Gaussian, (C) Mexican hat, and (D) dual-stage.

**Figure 5.**
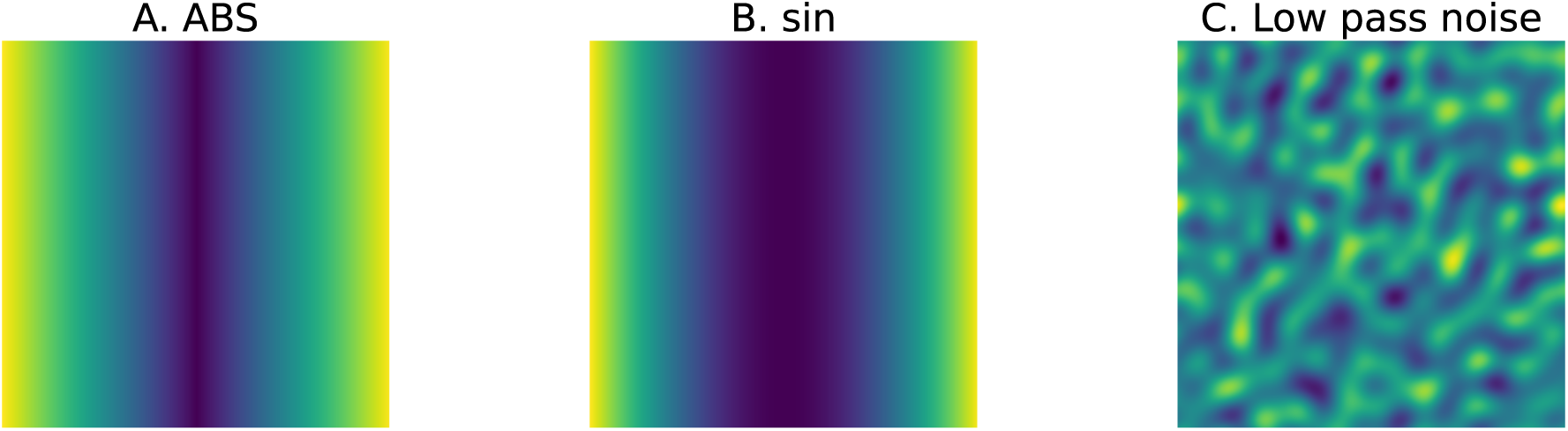
Spatially inhomogeneous parameter (*F*_base_) definition. (A) ABS (absolute value) parameter and (B) sine function parameters are used to limit the region of interdigitation, and (C) LowPass parameter is used to introduce stochasticity in the final pattern.

**Figure 6.**
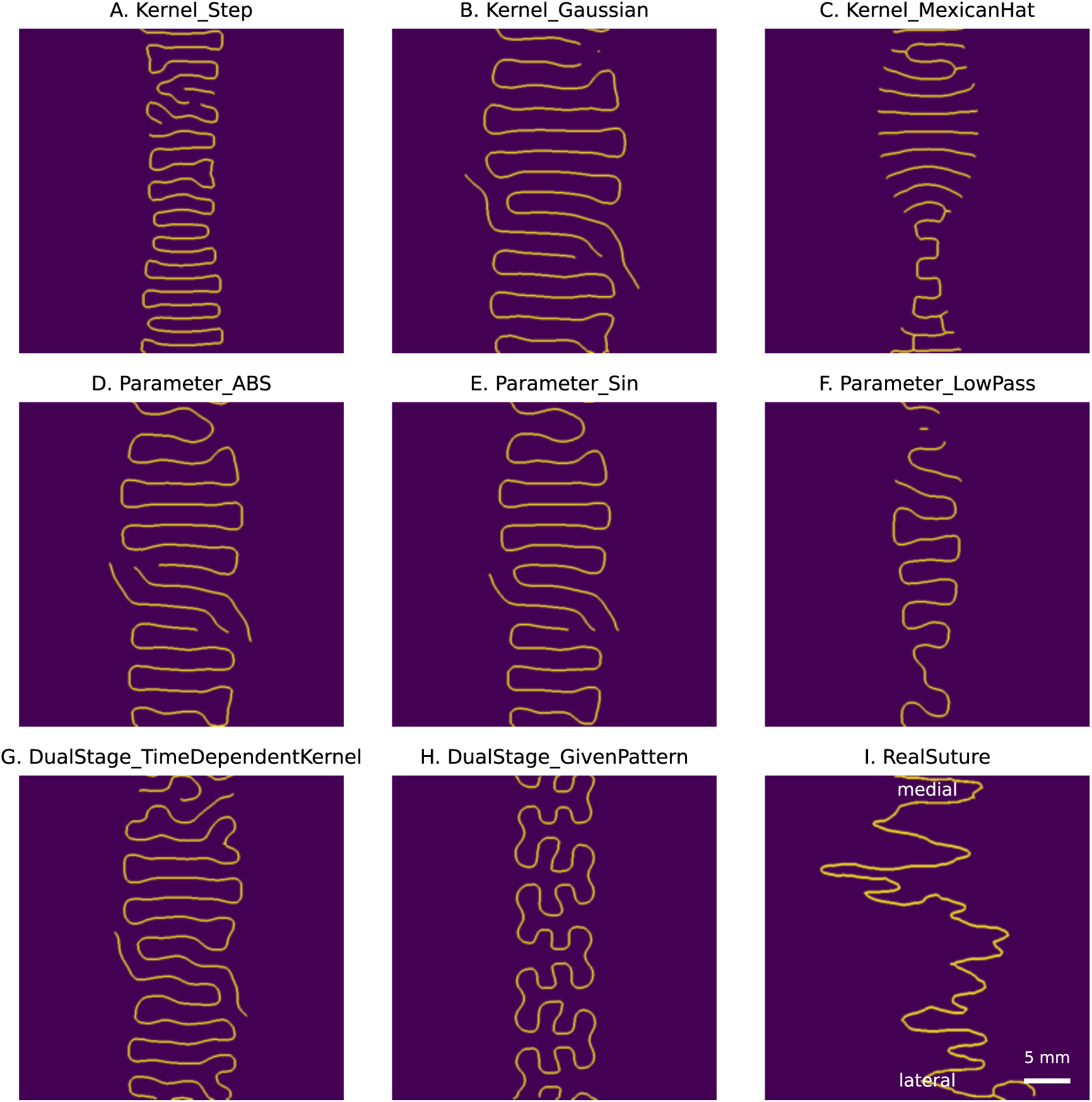
Skeletonized images of the patterns generated by mathematical models. (A) Step-function kernel. (B) Gaussian kernel. (C) Mexican hat kernel. (D) ABS-type *F*_base_. (E) Sine-type *F*_base_. (F) Low-pass noise. (G) Time-dependent kernel. (H) Pattern started from a sine curve. (I) Pattern of a real suture (top, medial; bottom, lateral).

**Figure 7.**
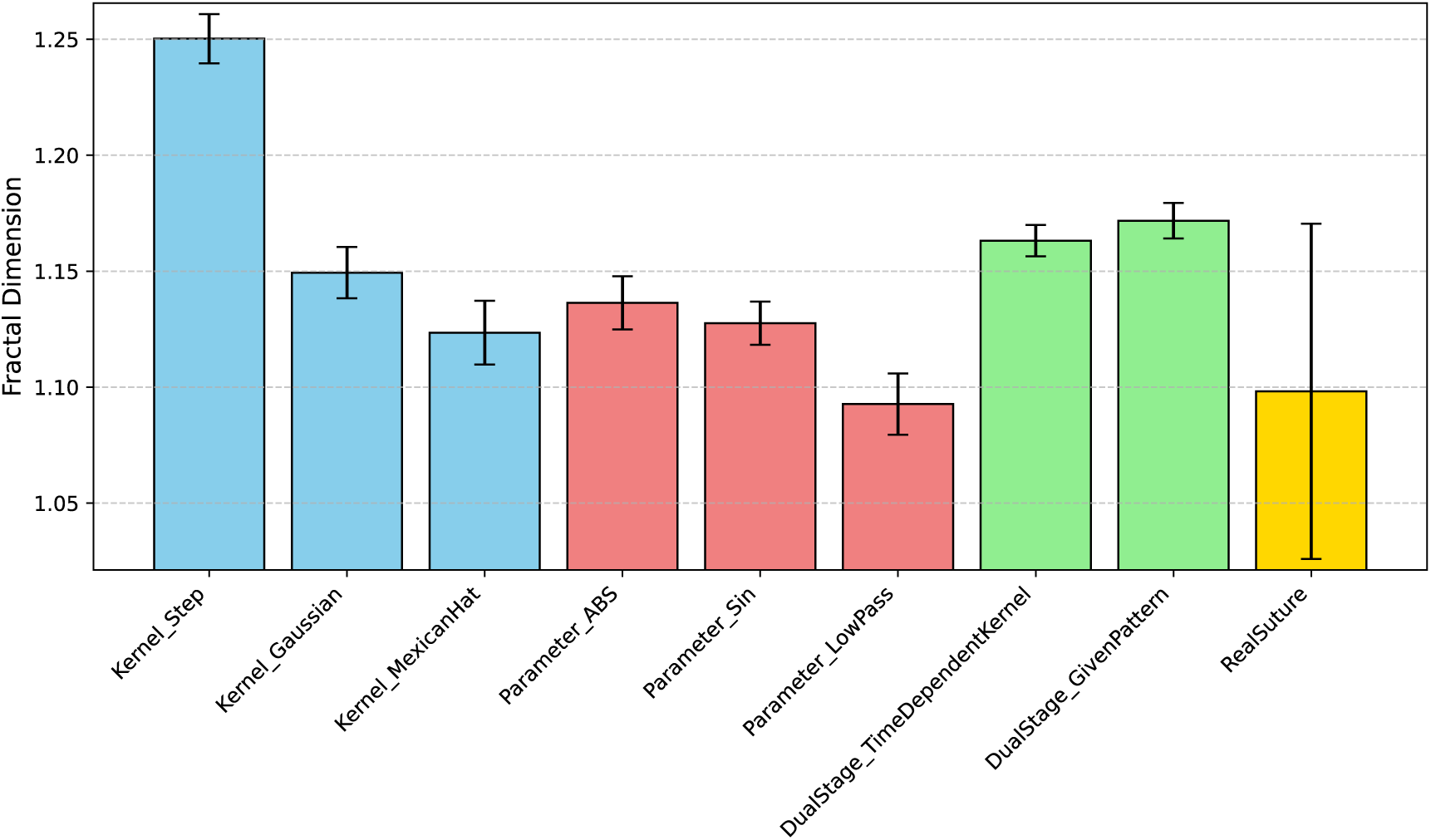
The fractal dimension of patterns generated by mathematical models. The error bars represent the standard deviation.

However, the observed FD increase is more plausibly explained by increased geometric curvature at specific spatial scales, which raises box occupancy, rather than by robust self-similar scaling across scales (Supporting Information, Section S2; Fig. S1). In this context, higher FD should be interpreted as increased local geometric complexity, not as direct evidence of clear self-similarity. This interpretation is also consistent with two limitations of the current setting: noise was added only to the initial condition, and its amplitude range was modest. Although the Step-kernel model produced relatively high FD values, likely due to its broad frequency content, FD alone cannot represent the full morphological similarity to real sutures.

Agreement in FD should be distinguished from agreement in morphology and, ultimately, from support for a model’s underlying biological mechanism. Similar FD values can accompany visually similar morphologies, as in models E and F, but can also arise from apparently distinct morphologies, as in models G and H (Fig. 6). Future work should combine FD with additional shape descriptors to evaluate how well models reproduce suture morphology more comprehensively. Consistent with comparative assessments of suture complexity metrics (White et al., 2020), FD should ideally be combined with complementary descriptors such as sinuosity and spectral measures when the goal is classification or broad morphometric comparison. Furthermore, morphological agreement alone does not validate a model’s underlying biological mechanism, because different types of models can generate similar patterns. Biological interpretation therefore requires evaluating both morphological agreement and independent evidence for the processes represented in each model.

### 4.4 Finite-Range Fractal-Like Nature of the Suture Line Model

In the suture line model developed in this study, the FD was nearly 1 at scales smaller than half the wavelength and approached 2 at larger scales (Fig. 3). Consequently, it was evaluated as an intermediate value overall. This is thought to be because, at the microscopic scale, the structure behaves as a one-dimensional curve, whereas at larger scales, the entire wave-like structure is reflected as a region with two-dimensional extent. Therefore, the box-counting dimension obtained in this study is not uniformly defined across a wide range of scales but is interpreted as reflecting different structural properties depending on the observation scale (Falconer, 2003).

Generally, known fractal structures include geometric and deterministic self-similar structures, statistical self-affine structures, statistical self-similar structures, multifractal structures, and fractal structures derived from dynamical systems (Falconer, 2003). However, the structure generated in this study could not be clearly classified into any of these categories, at least within the range observed in this study. The structure generated by the suture line model was not a fractal in the strict sense, but rather a “finite-range fractal-like structure” that exhibits fractal behavior only within a finite range of scales (Fig. 8, 9).

**Figure 8.**
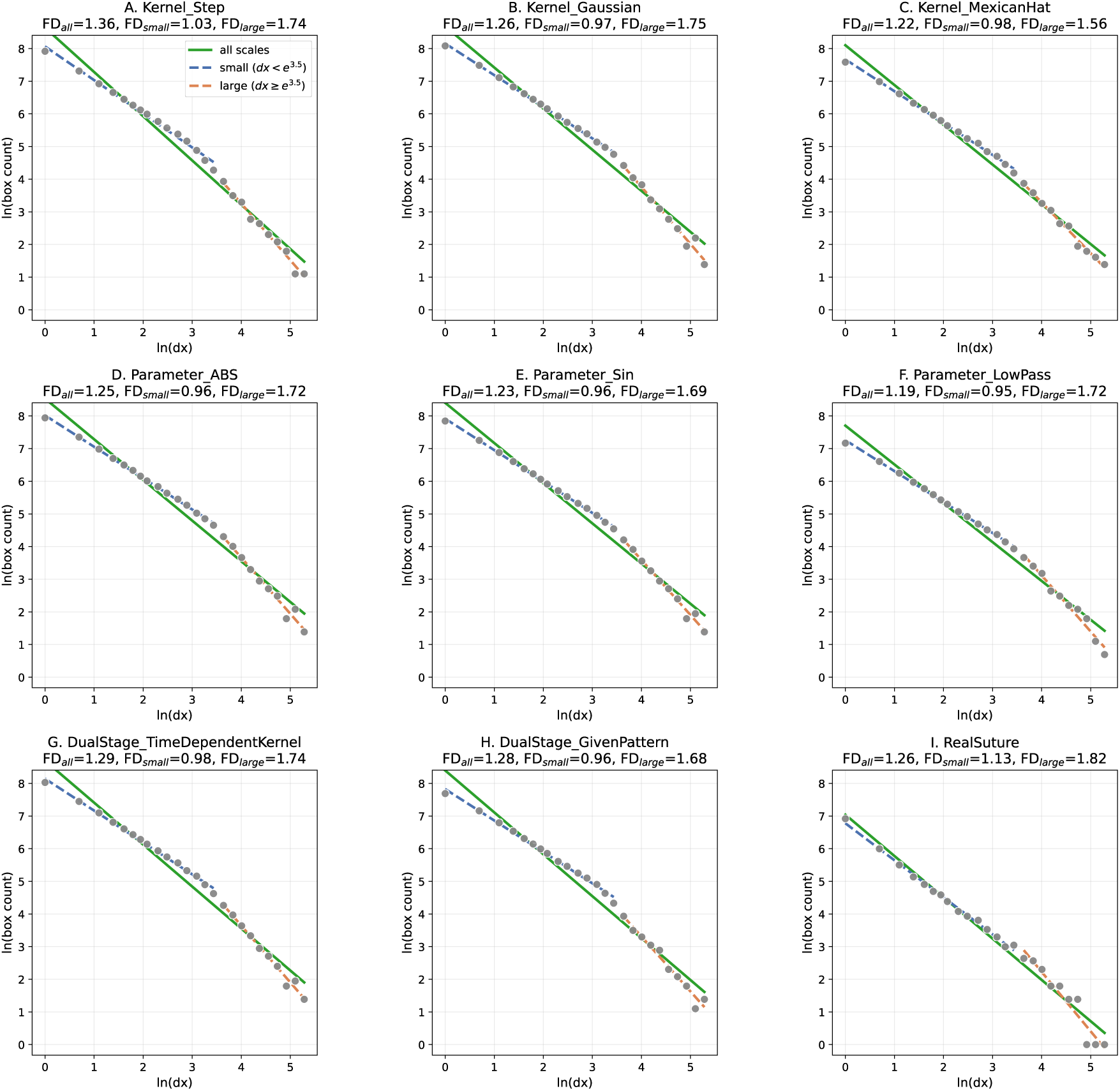
Log–log box-counting plots for representative skeletonized patterns from each model and an actual cranial suture. Points show occupied box counts at each scale, and solid green lines indicate linear fits for all scales used to estimate fractal dimension as the negative slope. Blue lines indicate linear fits for small scales, and orange lines indicate linear fits for large scales.

**Figure 9.**
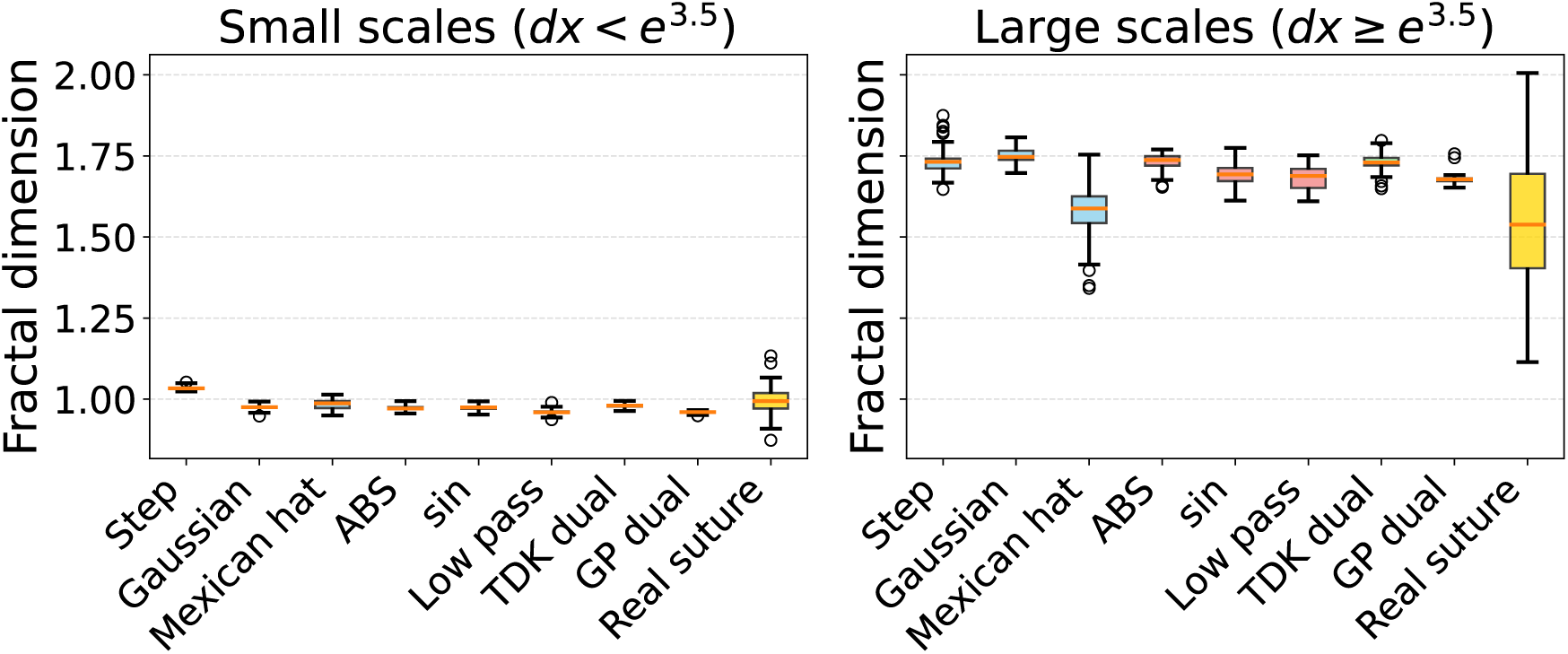
Differences in FD between large and small scales. In all data, including experimental observations, the small-scale dimension is close to 1, whereas the large-scale dimension is much larger than 1.

### 4.5 Possible mechanism for suture structure generation

Although several models produced FD values close to real data, resolving the causal biological and physical mechanisms responsible for actual suture morphology is beyond the scope of this paper. Here, we assess model–data agreement only at the level of FD under matched preprocessing and scale-range criteria. Candidate explanations discussed in prior work include multiscale structure emerging at different stages (Miura et al., 2009) and stochastic growth mechanisms such as the EW framework (Naroda et al., 2020). Discriminating among these mechanisms would require analysis beyond FD, which we plan to address in future studies.

### 4.6 Limitations

This study has several limitations. First, model–data agreement was evaluated primarily using FD, and therefore does not establish full morphological or developmental equivalence between simulated and actual sutures. Second, the simulations were performed over limited spatial and temporal scales, and longer-term growth dynamics remain to be examined. Third, all model variants shared a common set of initial conditions (Supporting Information Section S3), which suppresses between-replicate variability and likely accounts for the much narrower FD distributions produced by models compared with real specimens. Future work should explore how varying initial conditions, noise level, and noise timing affect the spread of FD across replicates.

### 4.7 Practical recommendations

In practical terms, we recommend that FD values be compared primarily within studies that share the same image-processing workflow, box-counting implementation, skeletonization status, and scale range. Cross-study or meta-analytic syntheses of published FD values should be undertaken only when these measurement conditions are matched or reported in sufficient detail to allow harmonization; otherwise, apparent differences may reflect methodological rather than biological variation. Ideally, sharing morphological data with calibration can enable accurate cross-study meta-analysis.

### 4.8 Future directions

Machine learning progress makes it feasible to compare patterns in latent space and build automated pipelines for morphological classification using unsupervised approaches or foundation models (Hishinuma et al., 2025). In addition, scale selection and preprocessing should be treated as part of the measurement protocol rather than merely as sources of error. Because the effective observation regime changes with box size (approximately half the wavelength as a practical boundary), FD reflects predominantly one-dimensional curve geometry at small scales, while larger scales capture more two-dimensional spatial organization. This suggests that using multiple FD values obtained under different pre-processing and scale settings can provide complementary FD-based features for pattern comparison and classification, beyond what a single FD value can capture.

## 5. Acknowledgments

This work is financially supported by JSPS Kakenhi (24K02036).

## 7. Data availability statement

The phase-field simulation code is available at https://github.com/haiken-hub/SutureModel. The fractal-dimension estimation library (box-counting) is available at https://github.com/haiken-hub/ScalingFractalTools.

## Supporting Information

### S1 Fractal geometry

Fractal dimension is a general term for metrics that measure the complexity of structure (Falconer, 2003). Generally, fractal dimension refers to the Hausdorff dimension. In a broader sense, box dimension and compass dimension—approximations of the Hausdorff dimension using methods such as the box-counting method or the divider method— are also included. Furthermore, because the required *&*-covering method differs in their derivation processes, the values for each method do not necessarily remain the same (Falconer, 2003).

The Hausdorff dimension was introduced by Felix Hausdorff (Hausdorff, 1918). For the Koch curve, one of the figures constructed by a recursive procedure, the Hausdorff dimension is approximately 1.26 (Mandelbrot, 1982). The Hausdorff dimension of a fractal structure assumes a non-integer value and is thereby distinguished from non-fractal structures, whose Hausdorff dimensions take integer values, such as a straight line with a Hausdorff dimension 1 or a plane with a Hausdorff dimension 2 (Mandelbrot, 1982).

In the box-counting method, the box dimension is derived by considering a covering using square subdivisions instead of the *&*-covering used in the derivation of the Hausdorff dimension. The compass dimension in the divider method is derived by considering circle coverings. For completely self-similar figures, these coverings coincide with the most efficient covering used for the Hausdorff dimension, so both dimensions take the same value (Falconer, 2003).

### S2 Box-counting dimension of sine curve

To understand why the two-phase box-counting dimension described above arises, we measured the box-counting dimension of a sine wave, which does not fit the classical definition of a fractal structure, and observed its characteristics. While the slope is 1 at small spatial scales, a dimension greater than 1 is observed when the box size is set to approximately half the wavelength of the wave. This can be interpreted as the box-counting dimension increasing because, at spatial resolutions exceeding this scale, the sine wave appears as a two-dimensional rectangle.

**Figure S1.**
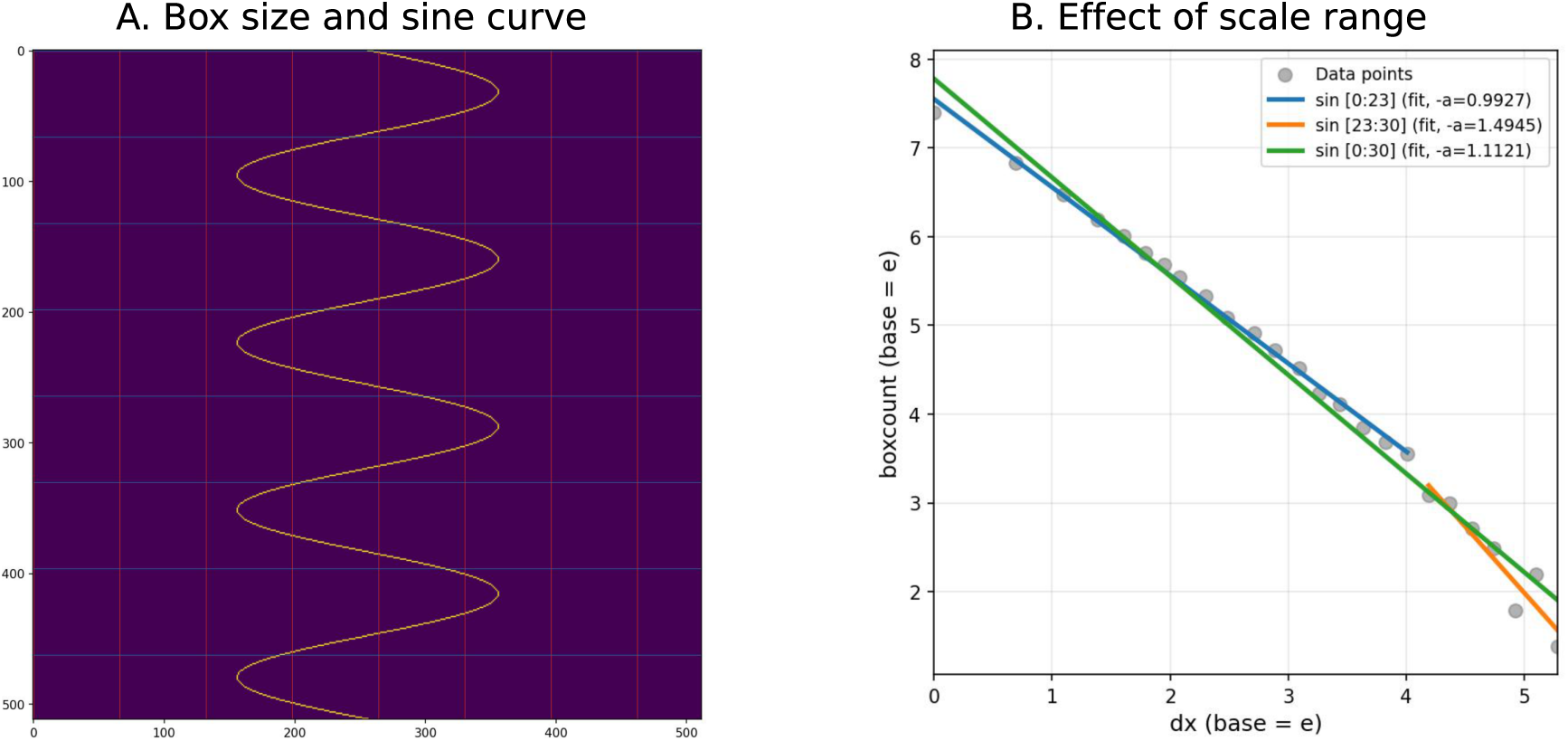
The relationship between fractal dimension and the amplitude and wavelength of a sine wave. (A) Scale selection effect. (B) The relationship between the box size at which the fractal dimension changes and the sine wave.

### S3 Detail of the model description

#### Base model

We utilized a mathematical model based on the interface equation (Yoshimura et al., 2016). We used a straight band-like region with a small interface perturbation. We defined the interface growth speed *V* as dependent on local interface curvature *∋* and nonlocal interaction kernel *K*. The interface equation is numerically implemented with the Allen-Cahn equation, in which bone is represented as *U* = 0 and mesenchyme is represented as *U* = 1.

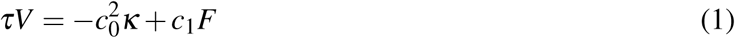

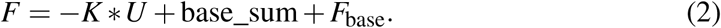

The initial band used as the signal reference was defined by *u*(**x**, 0)= 1 where (*y -n/*2)^2^ *<* (*x*_0_*/dx/*2)^2^ and 0 otherwise, with small integer perturbations (*±*10 grid points) on the band edge. With *n* = 512, *x*_0_ = 5.0, and *dx* = 0.5, the reference half-width is 20 grid points, and the band is aligned along the *x*-direction. The signal reference base_sum is computed from the unperturbed reference band *u*_0_ and the kernel using the same convolution convention as *K* \**U*. Thus, the threshold term in the original interface notation is not an independently specified model parameter; it is determined by the initial band width and the kernel *K*.

The interface equation (1) is implemented using the phase field as follows:

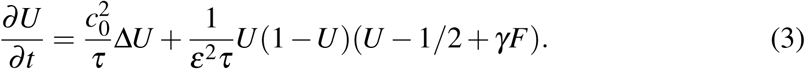

Here *%* is the coupling coefficient between the growth-factor signal *F* and the phase-field reaction term. It is not an independently fitted parameter but follows from matching the interface equation (1):

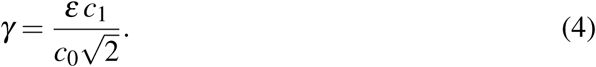

With *Ɛ* = 1, *c*_0_ = 0.33, and *c*_1_ = 0.11 (Table 2), *γ ≃* 0.236. Biologically, *γ* controls how strongly local growth-factor signalling (through *F*) promotes osteogenesis (*U* 1) versus suture maintenance (*U* 0) at the interface; larger *c*_1_ or smaller *c*_0_ yields larger *%* and a stronger morphogenetic response.

*Ɛ* is a free parameter that determines the interface thickness of the phase field; we set *Ɛ* = 1.0 in all simulations. Shared model parameters are summarized in Table 2, and model-variant settings are described in the model-specific subsections below.

In addition, we expanded this model as follows to reproduce patterns with a fractal dimension *FD >* 1.

#### Kernel design

##### Step Kernel

The step kernel assumes that spatial interaction is constant up to a certain length *l*_1_. In the `Kernel_Step` model, the uniform disk kernel used amplitude *w*_1_ = 0.006 and radius *l*_1_ = 13.8 in grid units. We used *t* = 936 snapshot for FD analysis.

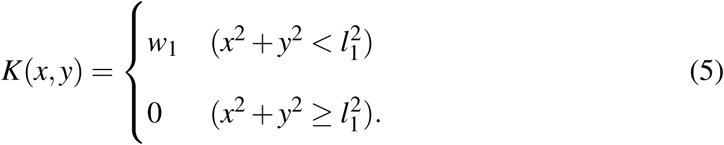

##### Gaussian Kernel

The Gaussian kernel assumes that spatial interaction depends on distance as follows:

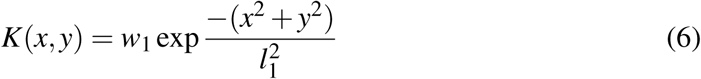

The `Kernel_Gaussian` model used *w*_1_ = 0.006, *l*_1_ = 8, *w*_2_ = 0, and *l*_2_ = 1 (*w*_2_ and *l*_2_ are unused in the single-Gaussian kernel). We used *t* = 2496 snapshot for FD analysis.

##### Mexican Hat Kernel

We use the Mexican hat kernel, assuming short-range stimulation and long-range inhibition.

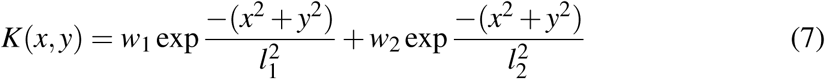

The `Kernel_MexicanHat` model used *w*_1_ = 0.00708, *l*_1_ = 8, *w*_2_ = *-*0.00118, and *l*_2_ = 16. We used *t* = 936 snapshot for FD analysis.

#### Spatially inhomogeneous parameters

##### Regional inhibition

We define two types of inhibitor distribution functions (scaled by 0.1 in all simulations):

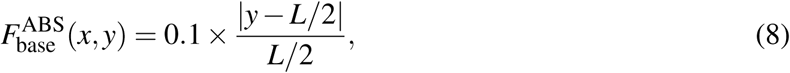

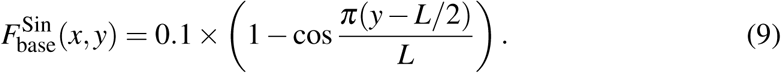

The initial osteogenic band lies near *y* = *L/*2; *F*_base_ depends on *y* only and modulates osteogenic tendency across the field width. The ABS model (`Parameter_ABS`) used the Gaussian kernel (*w*_1_ = 0.006, *l*_1_ = 8) with 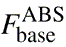. We used *t* = 1976 snapshot for FD analysis. The Sin model (`Parameter_Sin`) used the same Gaussian kernel with 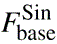. We used *t* = 1976 snapshot for FD analysis.

##### Low-frequency noise

To introduce stochasticity to the system, we added low-frequency noise *∗*_low_(*x*) (low-pass filtered noise) to *F*_base_. We defined the low-frequency noise as follows:

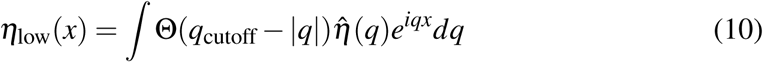

O(*x*) represents the Heaviside function, *q* denotes wave number (distinct from both the curvature *∋* in Eq. 1), and *q*_cutoff_ = 0.02 is the wave-number cutoff (in units of 1*/*pixel). The Low-pass model (`Parameter_LowPass`) used the Gaussian kernel (*w*_1_ = 0.006, *l*_1_ = 8) with low-pass noise scaled by 0.1. We used *t* = 1040 snapshot for FD analysis

#### Time-dependent parameters

##### Time-dependent kernel (TDK) — two-stage pipeline

The TDK model used two separate Allen–Cahn runs rather than a runtime kernel switch. Stage 1 ran the Gaussian kernel (*w*_1_ = 0.006, *l*_1_ = 8) for the standard 10,000 steps and stored snapshots at regular intervals. The snapshot at *T*_0_ = 1456 was extracted and used as the initial condition *u*_0_ for stage 2. Stage 2 ran an independent Allen–Cahn integration with a modified kernel (*w^⇔^*_1_ = 0.007, *l*_1_*^⇔^* = 6) for an additional 10,000 steps (*T*_1_ = 2500 in stage-2 time), corresponding to the equations

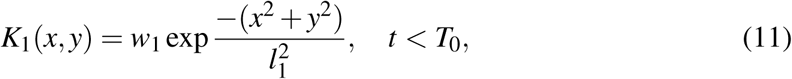

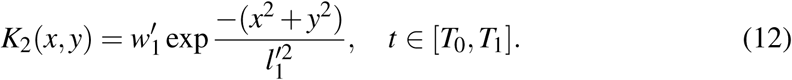

The FD analysis for DualStage_TDK used snapshot of *t* = 728 in stage-2 time.

##### Approximation of early-stage pattern

We describe the pattern at *T*_0_ as follows (sine varies along *x*; the band is bounded in *y*):

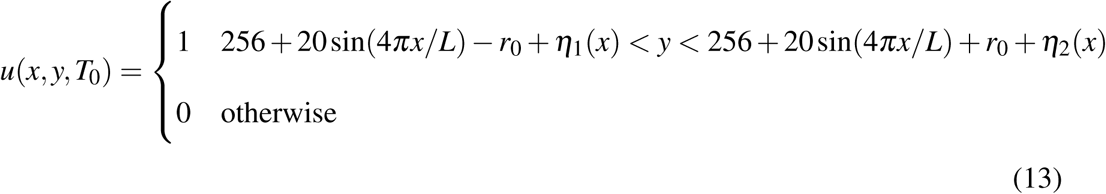

where *r*_0_ = 20 (half-width of the band in grid units) and *∗_i_*(*x*) represents uniform integer noise on the interval (*-*3, 3) drawn independently at each *x*. The `DualStage_GP` model used this sine-band initial condition with the stage-2 kernel *w*_1_ = 0.007 and *l*_1_ = 6. The FD analysis used snapshot at *t* = 1040.

#### Numerical simulation

Numerical simulations were performed on a two-dimensional uniform grid of 512 *×* 512 points with a spatial step of *dx* = 0.5, using a split-step semi-implicit scheme (explicit update for the nonlinear/nonlocal reaction terms and FFT-based implicit update for diffusion). The time step was fixed at *dt* = 0.25, and each integration segment ran for a fixed 10,000 steps (*T* = 2500 in simulation time), without an additional convergence-based stopping rule. Since the interface instability varies among models, the snapshot for FD analysis was chosen so that suture occupies about 1/3 of the total area. Because diffusion was advanced in Fourier space, the implementation corresponds to periodic boundary conditions. Initial conditions were generated as binary fields with a noisy central band and then shared across model variants via a common initial-condition set, so between-model comparisons used matched starts. A seed value of 32 is passed in the solver interface; however, when precomputed initial fields are supplied (the standard workflow), stochasticity is determined by the initial-condition generation step. The number of replicates is set by the number of available input samples (45 in the current dataset), with an optional cap through `MAX_SAMPLES` for reduced runs.

Each model variant exported the phase-field snapshot at the model-specific index *k* stated in the corresponding model description above; these snapshots were not necessarily from the final step of the integration. Snapshot times approximate *t ≃* 104 *k* (derived from 416 *× dt* = 416 × 0.25, where 416 is the number of steps between stored outputs, `steps_per_output`). For the `DualStage_TDK` model, the FD snapshot is at *t ≃* 728 within stage 2, while *T*_1_ = 2500 denotes the end of stage-2 integration.

## References

Buezas, G., Becerra, F., & Vassallo, A. (2017). Cranial suture complexity in caviomorph rodents (*Rodentia; Ctenohystrica*). J. Morphol., 278, 1125–1136. 10.1002/jmor.20699.

Byron, C. D. (2009). Cranial suture morphology and its relationship to diet in *Cebus*. J. Hum. Evol., 57, 649–655. 10.1016/j.jhevol.2008.11.006.

Cohen, M. M., & MacLean, R. E. (2000). Craniosynostosis. Diagnosis, Evaluation, and Management. London, England: Oxford University Press.

Costamagna, D., Cassini, G., Cabral, V., Schmidt, G. I., & Ferrero, B. S. (2026). Fractal dimension of dental crowns: An innovative method for age estimation in toxodonts. J. Exp. Zool. B, 346, 200–213. 10.1002/jez.b.23319.

Darawsheh, A. F., Kolarovszki, B., Hong, D. H., Farkas, N., Taheri, S., & Frank, D. (2023). Applicability of fractal analysis for quantitative evaluation of midpalatal suture maturation. J. Clin. Med., 12, 4189. 10.3390/jcm12134189.

Di Guida, N. S., & Cassini, G. H. (2026). Fractal dimension and suture complexity during postnatal ontogeny in neotropical deer in relation to sexual dimorphism and other biological features. J. Exp. Zool. B, 346, 172–187. 10.1002/jez.b.23300.

Drake, R. L., Moses, K., Vogl, A. W., & Mitchell, A. W. M. (2014). Gray’s Anatomy for Students. Elsevier, third edition.

Dutch, S. I. (1993). Linear Richardson plots from non-fractal data sets. Math. Geol., 25, 737–751. 10.1007/BF00893176.

Eden, M. (1961). A two-dimensional growth process. In: Fourth Berkeley Symposium on Mathematical Statistics and Probability, volume 4, pages 223–239.

Falconer, K. (2003). Fractal Geometry. New York, NY: John Wiley & Sons Ltd., second edition.

Gibert, J., & Palmqvist, P. (1995). Fractal analysis of the Orce skull sutures. J. Hum. Evol., 28, 561–575. 10.1006/jhev.1995.1042.

Górski, A. Z., & Skrzat, J. (2006). Error estimation of the fractal dimension measurements of cranial sutures. J. Anat., 208, 353–359. 10.1111/j.1469-7580.2006.00529.x.

Hartwig, W. C. (1991). Fractal analysis of sagittal suture morphology. J. Morphol., 210, 289–298. 10.1002/jmor.1052100307.

Hausdorff, F. (1918). Dimension und äußeres Maß. Math. Ann., 79, 157–179. 10.1007/BF01457179.

Herring, S. W. (2008). Mechanical influences on suture development and patency. Front. Oral Biol., 12, 41–56.

Hishinuma, H., Takigawa-Imamura, H., & Miura, T. (2025). Data-driven discovery and parameter estimation of mathematical models in biological pattern formation. PLoS Comput. Biol., 21, e1012689. 10.1371/journal.pcbi.1012689.

Klinkenberg, B. (1994). A review of methods used to determine the fractal dimension of linear features. Math. Geol., 26, 23–46. 10.1007/bf02065874.

Long, C. A. (1985). Intricate sutures as fractal curves. J. Morphol., 185, 285–295. 10.1002/jmor.1051850303.

Long, C. A., & Long, J. E. (1992). Fractal dimensions of cranial sutures and waveforms. Acta Anat. (Basel), 145, 201–206. 10.1159/000147366.

Lynnerup, N., & Jacobsen, J. C. B. (2003). Brief communication: Age and fractal dimensions of human sagittal and coronal sutures. Am. J. Phys. Anthropol., 121, 332–336. 10.1002/ajpa.10260.

Mandelbrot, B. (1982). The Fractal Geometry of Nature. London: Times Books.

Masuda, Y., & Yohro, T. (1987). Are there any regularities in cranial sutures? Okajimas Folia Anat. Jpn., 64, 39–45.

Miura, T., Perlyn, C. A., Kinboshi, M., Ogihara, N., Kobayashi-Miura, M., Morriss-Kay, G. M., & Shiota, K. (2009). Mechanism of skull suture maintenance and interdigitation. J. Anat., 215, 642–655. 10.1111/j.1469-7580.2009.01148.x.

Monteiro, L. R., & Lessa, L. G. (2000). Comparative analysis of cranial suture complexity in the genus *Caiman* (*Crocodylia*, Alligatoridae). Braz. J. Biol., 60, 689–694. 10.1590/S0034-71082000000400021.

Naroda, Y., Endo, Y., Yoshimura, K., Ishii, H., Ei, S.-I., & Miura, T. (2020). Noise-induced scaling in skull suture interdigitation. PLoS One, 15, e0235802. 10.1371/journal.pone.0235802.

Nicolay, C. W., & Vaders, M. J. (2006). Cranial suture complexity in white-tailed deer (*Odocoileus virginianus*). J. Morphol., 267, 841–849. 10.1002/jmor.10445.

Ok, U., & Kaya, T. U. (2022). Fractal perspective on the rapid maxillary expansion treatment; evaluation of the relationship between midpalatal suture opening and dental effects. J. Stomatol. Oral Maxillofac. Surg., 123, 422–428. 10.1016/j.jormas.2021.09.002.

Oota, Y., Nagamine, T., Ono, K., & Miyazima, S. (2004). A two-dimensional model for sagittal suture of cranium. Forma, 19, 197–205.

Oota, Y., Ono, K., & Miyazima, S. (2006). 3D modeling for sagittal suture. Physica A, 359, 538–546. 10.1016/j.physa.2005.05.095.

Otsu, N. (1979). A threshold selection method from gray-level histograms. IEEE Trans. Syst. Man Cybern., 9, 62–66. 10.1109/TSMC.1979.4310076.

Skrzat, J., Walocha, J., & Zawiliński, J. (2004). A note on the morphology of the metopic suture in the human skull. Folia Morphol. (Warsz.), 63, 481–484.

Twigg, S. R. F., & Wilkie, A. O. M. (2015). A genetic-pathophysiological framework for craniosynostosis. Am. J. Hum. Genet., 97, 359–377. 10.1016/j.ajhg.2015.07.006.

Van Rossum, G., & Drake, F. L. (2009). Python 3 Reference Manual. Scotts Valley, CA: CreateSpace.

White, H. E., Camaiti, M., Tucker, A. S., Watanabe, A., & Goswami, A. (2026). A suture in time: The ontogeny of cranial suture morphology in mammals. J. Anat., 248, 501–516. 10.1111/joa.70035.

White, H. E., Clavel, J., Tucker, A. S., & Goswami, A. (2020). A comparison of metrics for quantifying cranial suture complexity. J. R. Soc. Interface, 17, 20200476. 10.1098/rsif.2020.0476.

Yoshimura, K., Kobayashi, R., Ohmura, T., Kajimoto, Y., & Miura, T. (2016). A new mathematical model for pattern formation by cranial sutures. J. Theor. Biol., 408, 66–74. 10.1016/j.jtbi.2016.08.003.

Yu, J. C., Wright, R. L., Williamson, M. A., Braselton, J. P., & Abell, M. L. (2003). A fractal analysis of human cranial sutures. Cleft Palate Craniofac. J., 40, 409–415.

Zollikofer, C. P. E., & Weissmann, J. D. (2011). A bidirectional interface growth model for cranial interosseous suture morphogenesis. J. Anat., 219, 100–114. 10.1111/j.1469-7580.2011.01386.x.

